# Wavelet Phase Coherence of Ictal Scalp EEG-Extracted Muscle Activity (SMA) as a Biomarker for Sudden Unexpected Death in Epilepsy (SUDEP)

**DOI:** 10.1101/2024.02.04.578837

**Authors:** Adam C. Gravitis, Krishram Sivendiran, Uilki Tufa, Katherine Zukotynski, Yotin Chinvarun, Orrin Devinsky, Richard Wennberg, Peter L. Carlen, Berj L. Bardakjian

**Author notes:** Co-first author.

## Abstract

**Objective:** Approximately 50 million people worldwide have epilepsy and 8-17% of the deaths in patients with epilepsy are attributed to sudden unexpected death in epilepsy (SUDEP). The goal of the present work was to establish a biomarker for SUDEP so that preventive treatment can be instituted.

**Approach:** Seizure activity in patients with SUDEP and non-SUDEP was analyzed, specifically, the scalp EEG extracted muscle activity (SMA) and the average wavelet phase coherence (WPC) during seizures was computed for two frequency ranges (1-12 Hz, 13-30 Hz) to identify differences between the two groups.

**Main results:** Ictal SMA in SUDEP patients showed a statistically higher average WPC value when compared to non-SUDEP patients for both frequency ranges. Area under curve for a cross-validated logistic classifier was 81%.

**Significance:** Average WPC of ictal SMA is a candidate biomarker for early detection of SUDEP.

## Introduction

Epilepsy is a common chronic neurological disorder characterized by recurrent seizures. Sudden unexpected death in epilepsy (SUDEP) occurs in approximately 1 in 1000 people with epilepsy each year [1] and typically occurs after convulsive seizures in sleep, followed by cardio-respiratory dysfunction and impaired arousal which may be caused by spreading depression or epileptiform activity involving the brainstem [1-6]. A biomarker for epilepsy patients at high SUDEP risk could enable earlier and more aggressive preventive interventions. Electromyography (EMG) can detect tonic-clonic seizures [7]. Scalp muscle activity (SMA) has been shown as a useful diagnostic for detection of tonic-clonic seizures, an established risk factor for SUDEP [8-9]. In our retrospective study, since limb EMG was not recorded, we extracted electrical muscle activity from scalp electrodes (i.e., SMA). Ictal SMA had an average wavelet phase coherence (WPC) in two frequency ranges that was significantly different in the SUDEP group as compared to the non-SUDEP group, identifying average WPC as a candidate biomarker for SUDEP.

## Method

### Data Acquisition

Scalp EEG recordings were obtained from 5 non-SUDEP and 7 definite SUDEP patients (sudden, unexpected death of a patient without relevant comorbidities, in which postmortem examination, including toxicology, does not reveal a cause of death other than epilepsy). Non-SUDEP controls were selected based on their similarity to SUDEP patients. EEG recordings were acquired using the Natus/Xltek EEG system with 19 or more electrodes. Although no recorded seizures were fatal, all SUDEP patients died within 3 years of their last available recordings. Patients categorized as non-SUDEP did not die within 10 years of their last available recordings.

Patients were undergoing presurgical evaluation in an epilepsy monitoring unit (EMU), with drug-resistant focal (temporal or extratemporal lobe) epilepsy and were not on anti-seizure medications at the time of recording. Apart from their definite SUDEP designation, the following data were not available in this retrospective study: simultaneous video EEG, sleep/wakefulness states, other medications, MRI findings, or non-epilepsy medical history.

The data were obtained through the consortium formed by the Toronto Western Hospital, the New York University (NYU) Comprehensive Epilepsy Center, and the Phramongkutklao Royal Army Hospital (Tables I and II). Ictal durations were identified from EEG scalp electrode recordings by board-certified neurologists/ electroencephalographers (Table 1).

**Table 1.** Ictal Patient Data (*one seizure from patient 9 was reserved for risk assessment; ^**^ one interictal segment from patient 11 was reserved for autocorrelation threshold selection, per Table 2).

| Patient | Classification | Age | Sex | Sampling Rate [Hz] | # of Ictal Recordings | Range of Ictal Durations [s] |
| --- | --- | --- | --- | --- | --- | --- |
| 1 | non-SUDEP | 28 | M | 500 | 2 | 73 - 126 |
| 2 | non-SUDEP | 19 | M | 512 | 3 | 60 - 138 |
| 3 | non-SUDEP | 42 | F | 512 | 1 | 86 |
| 4 | non-SUDEP | 40 | F | 512 | 1 | 69 |
| 5 | non-SUDEP | 28 | M | 512 | 5 | 54 - 109 |
| 6 | SUDEP | 13 | M | 256 | 2 | 76 - 84 |
| 7 | SUDEP | 21 | F | 256 | 1 | 241 |
| 8 | SUDEP | 43 | F | 512 | 6 | 62 - 272 |
| 9* | SUDEP | 47 | M | 200 | 3 | 110 - 474 |
| 10 | SUDEP | 30 | M | 256 | 1 | 117 |
| 11** | SUDEP | 47 | F | 500 | 1 | 63 |

### Protocol Approvals and Data Availability

The institutional review boards of the multi-centre consortium approved the study protocols and all patients gave informed consent. The anonymized datasets used in this study are available upon request. They are not publicly available due to institutional restrictions associated with original data acquisition protocols.

### SMA Extraction and Analysis

(1) Original EEG recordings ranged in sampling rate from 200 Hz to 512 Hz. All recordings were upsampled to 512 Hz, and low-pass filtered at 100 Hz. (2) EEG signals from each of the 19 standard electrodes of the international 10-20 system were decomposed into 30 components using singular spectrum analysis (SSA). (3) Notch filtering of 60 Hz and its harmonics. (4) Autocorrelation values for each SSA component were calculated. (5) An autocorrelation threshold was tuned to maximize EMG-like properties of extracted signal. (6) SSA components below the tuned autocorrelation threshold were extracted as EMG-like SMA signals. (7) WPC was calculated between retained components of each electrode pair. (8) Average values of WPC over ictal duration and 1-12 Hz and 13-30 Hz ranges were calculated. (9) Per-seizure spatial averages of WPC were derived from per-electrode temporal averages.

Reported differences between EEG and EMG signals [10] were reflected in the extracted SMA and retained EEG resulting from this methodology (Fig. 2). Extracted SMA, selected for lower autocorrelation values, had dominant power above 50 Hz, but also included activity in the 1-30 Hz range. Retained EEG was selected for higher autocorrelation and resulting in dominant power below 50 Hz.

**Figure 1.**
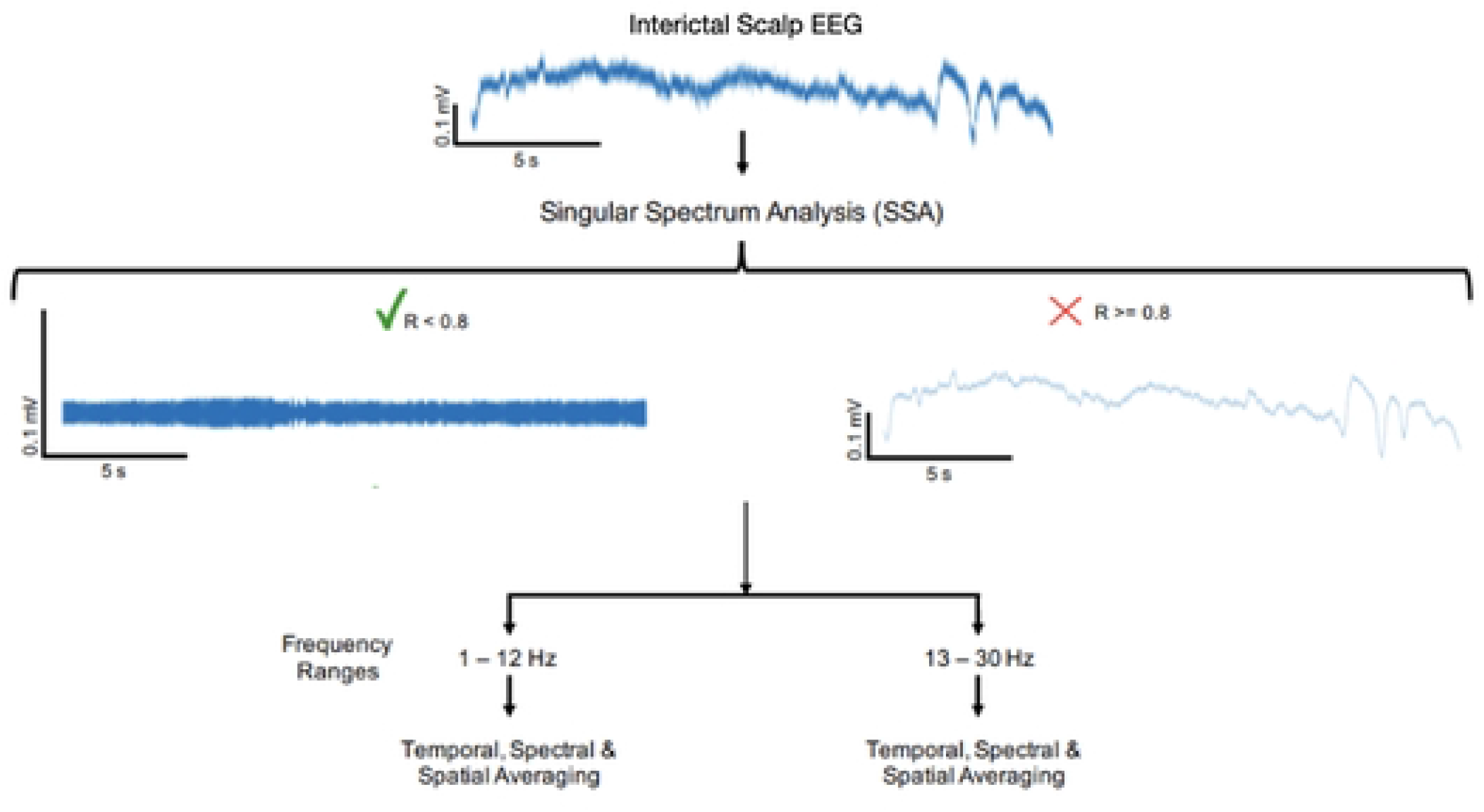
Block diagram showing methodology: the shown interictal EEG trace is from patient 12. Retained SSA components below the tuned autocorrelation threshold (R < 0.8) were summed to form the extracted SMA signal, which was analyzed in two frequency ranges: 1-12 Hz and 13-30 Hz.

**Figure 2.**
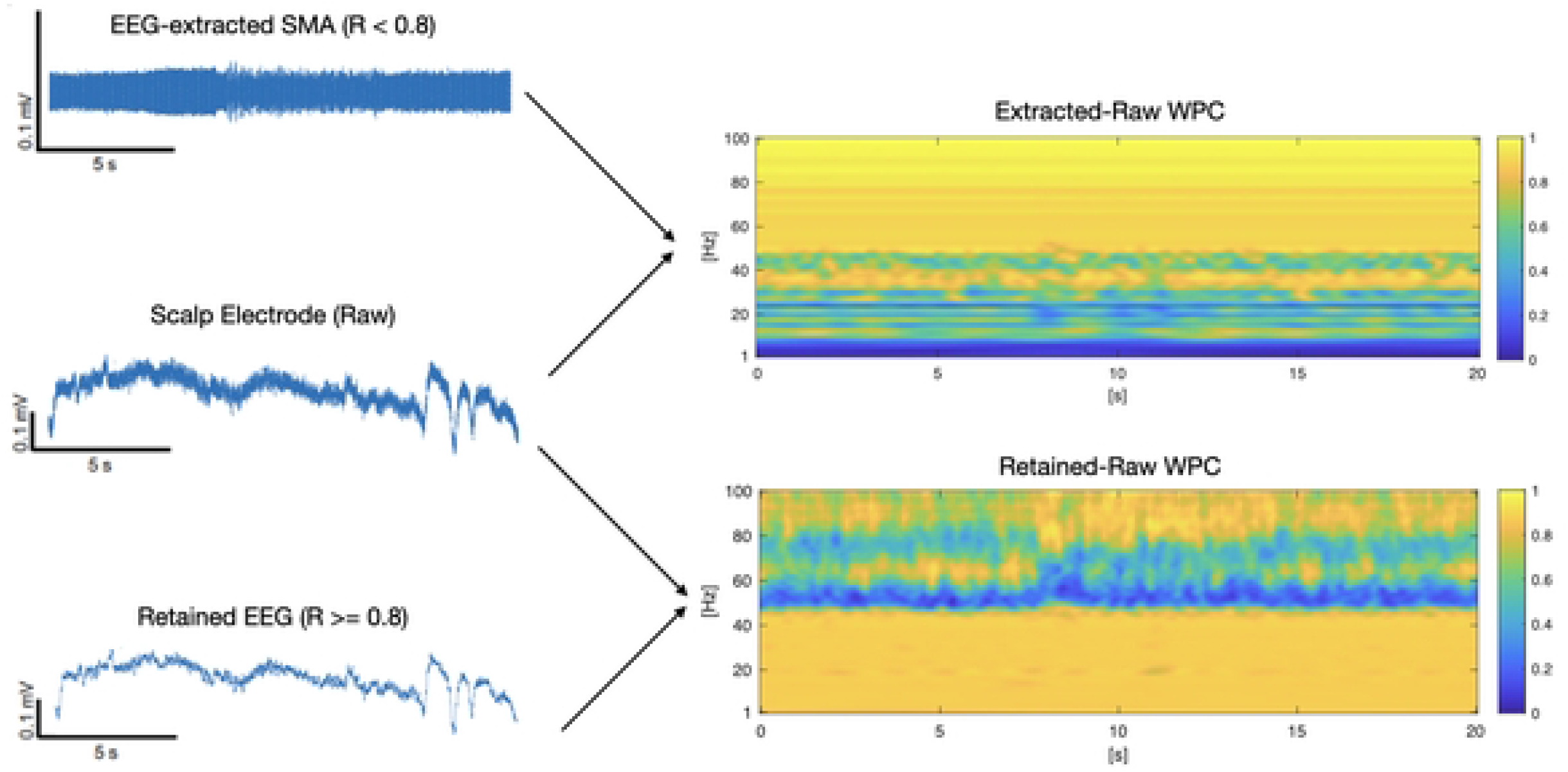
WPC of extracted SMA and retained EEG against raw electrode for patient 12.

### Singular Spectrum Analysis (SSA)

SSA using elementary grouping [10] was first performed on raw EEG data from ictal recordings to decompose the signal into its constituent components. This technique consists of first creating a trajectory matrix, T, from lagged versions of the time series x (in this case, a single EEG electrode recording). Next, the singular value decomposition of the trajectory matrix was taken. Using values obtained from this decomposition, the trajectory matrix was decomposed into a sum of L elementary matrices (matrices that have a rank of 1), where U and V were obtained from the singular value decomposition of the trajectory matrix, λ represents the eigenvalues of the trajectory matrix and k ranges from 0 to L-1.

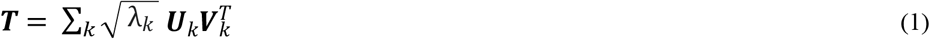

Finally, each of the L elementary matrices were hankelized and each resulting Hankel matrix was converted into a time series, where each diagonal value of the Hankel matrix corresponds to a sample in the time series.

### Autocorrelation Analysis

The autocorrelation of each SSA component was computed following equation (2) [10], where E is the expected value operation, s_1_(t) is the SSA component time series and s_2_ (t) = s_1_ (t-1).

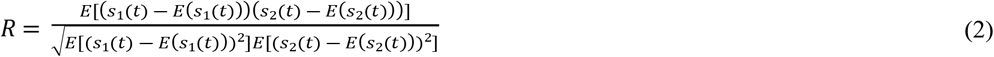

As muscle activity has a wide frequency range, it has a lower autocorrelation value than EEG signals. Reference [10] demonstrated an autocorrelation threshold can be used to differentiate an EEG signal from scalp muscle activity. We apply the technique to extract SMA from EEG.

The SSA components identified as EEG were removed and the remaining components summed to recover SMA.

### Autocorrelation Threshold Selection

Our analysis depended on an autocorrelation threshold value to distinguish extracted muscle activity from EEG rhythms (Fig. 3(b)). Autocorrelation thresholds between 0.5 and 0.95 were tested, at 0.05 intervals. Five inter-ictal EEG segments of SUDEP patients were used as controls, each spanning 62 s, and were processed to obtain resulting SMA signals for each threshold value. All control recordings were obtained from 2 SUDEP patients (2 from patient 11, 3 from patient 12) and were used for threshold tuning (Table 2).

**Table 2.**
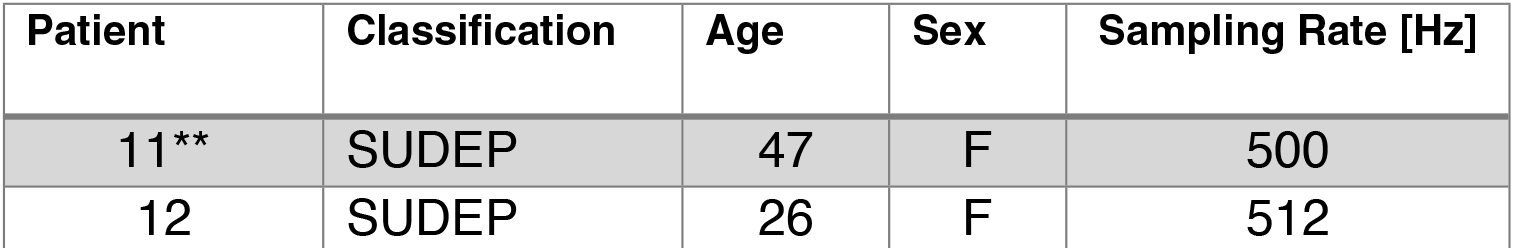
Interictal Patient Data, used for autocorrelation threshold selection.

**Figure 3.**
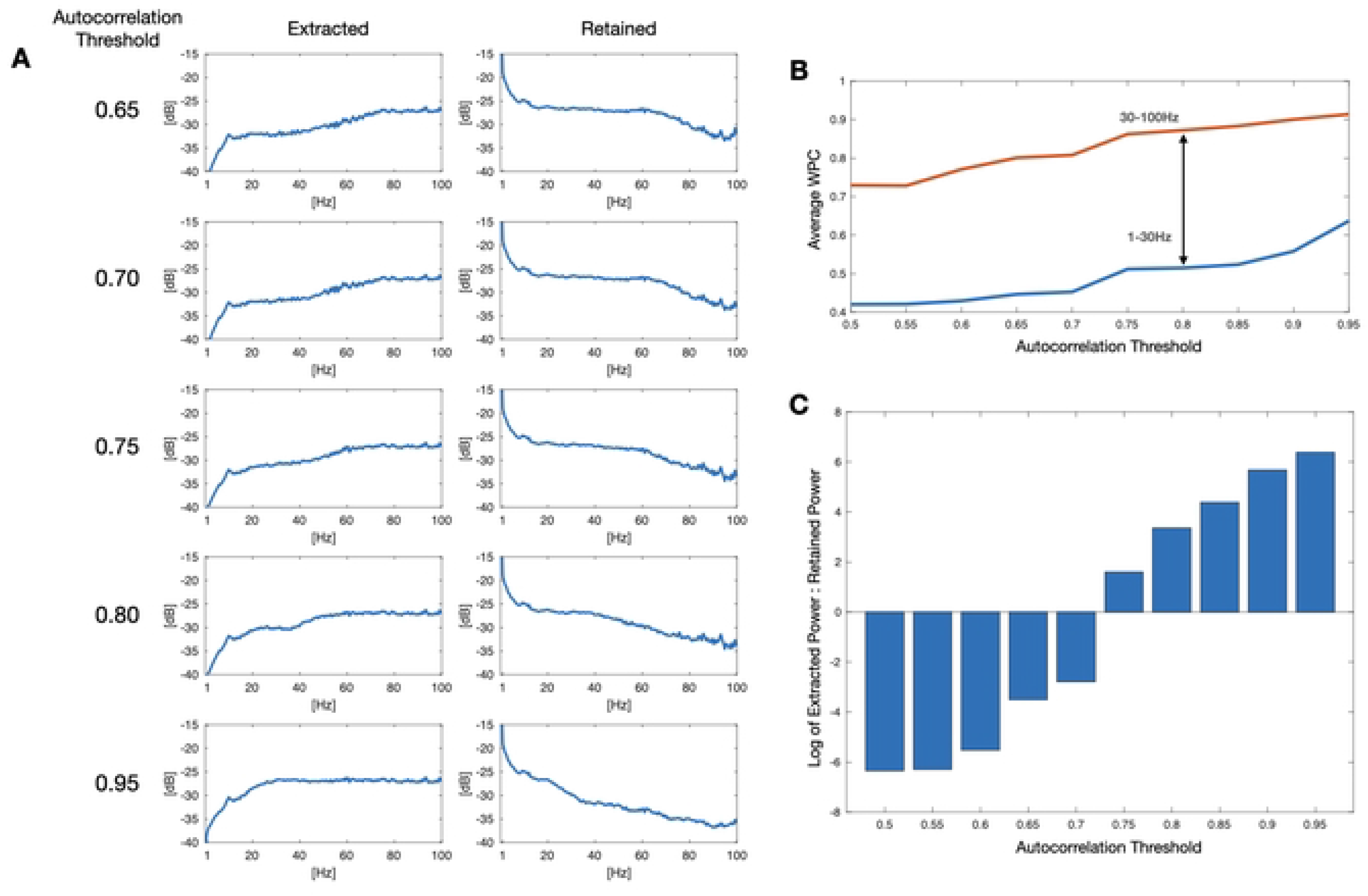
Autocorrelation threshold tuning. (A) Power spectra of extracted and retained signals for varying autocorrelation thresholds. (B) Comparison of average WPC between extracted SMA and raw electrode for 1-30 Hz and 30-100 Hz ranges: 0.75-0.85 maximized this difference. (C) The log of the ratio of extracted power between 50-70 Hz at different autocorrelation thresholds.

The WPC between each of the 5 resulting SMA signals and their corresponding scalp electrodes at each autocorrelation threshold. Electrodes FZ, FP1 and FP2 were selected due to the high presence of scalp muscle activity when compared to other electrodes, due to proximity of facial muscles. Each data matrix was averaged over time and averaged over two frequency ranges: 1-30 Hz and 31-100 Hz.

Fig. 3(b) shows the change in average WPC between extracted SMA and raw electrode for both 1-30 Hz and 31-100 Hz frequency ranges for varying autocorrelation thresholds. A threshold of 0.8 was selected as it maximized the difference between EMG-like retained SMA and retained EEG, based on power spectra (Fig. 3(a)), WPC (Fig. 3(b)), and ratio of power in the 50-70 Hz range (Fig. 3(c)).

Phase-phase cross-frequency coupling (PPC) analysis was performed between the extracted SMA and corresponding electrode for an ictal segment from patient 11 (Fig. 2), using an *n:m* PPC calculation [11-12]:

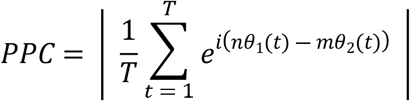

PPC analysis (Fig. 4(b)) confirmed strong coupling between the 50-70 Hz EMG-like frequencies of the raw electrode with lower frequencies of extracted SMA, most pronounced at 12-15 Hz.

**Figure 4.**
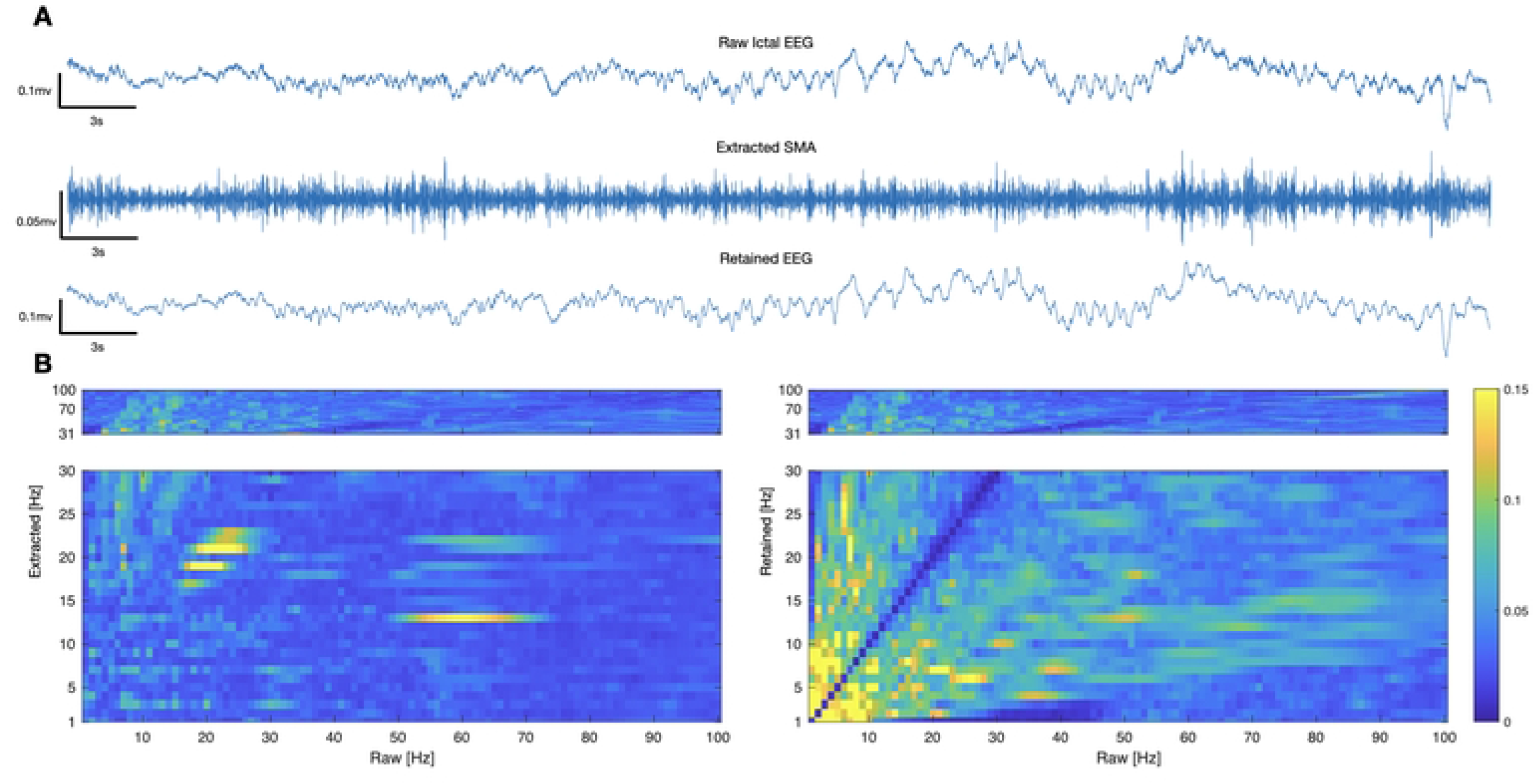
Phase coupling of extracted SMA with EMG signal in ictal segment of patient 12. (A) Raw ictal EEG, Extracted SMA, and Retained EEG. (B) Phase-phase cross-frequency coupling (PPC) between extracted SMA and raw electrode (left), and between retained EEG and raw electrode (right). Lower frequencies of extracted SMA are strongly coupled with higher frequencies of EMG.

### Wavelet Phase Coherence

The WPC of the SMA signals were obtained for each electrode pair. Since 19 electrodes from each ictal recording were used in the analysis, this resulted in 361 (19 x 19) WPC data matrices. For each data matrix, a time average was performed over the entire duration of the seizure. A frequency average was then performed over 1-12 Hz and 13-30 Hz.

Disregarding coherence entries between identical electrodes, each row of the resulting matrix was then averaged over the 18 column entries to obtain the average WPC on a per-electrode basis. To obtain the average WPC on a per seizure basis, each group of 19 electrodes was averaged.

### Estimation Statistics

Results are presented using estimation statistics as an alternative to null hypothesis significance testing [13].

### Risk Assessment

A logistic classifier was trained on the WPC values of both frequency bands of interest to produce a propensity score. One seizure from Patient 9 was withheld from training, in order to validate the risk assessment produced by the classifier.

## Results

### Validation of SMA Extraction

We hypothesized that the WPC of retained (following SMA removal) EEG would be distinct from that of the extracted SMA, as an initial validation of the extraction process.

Using the optimal autocorrelation threshold of 0.8, SMA was extracted from 5 SUDEP (12 seizure recordings) and 5 non-SUDEP (12 seizure recordings) patients. However, instead of discarding the SSA components which corresponded to an autocorrelation value greater than or equal to 0.8, the components were summed to obtain the retained EEG signal.

Next, average WPC was computed on a per electrode basis for the SMA and retained EEG signals. For both frequency ranges: 1-12 Hz and 13-30 Hz, Fig. 5(a) compares the average WPC for both signals for SUDEP patients and Fig. 5(b) compares the average WPC for both signals for non-SUDEP patients.

**Figure 5.**
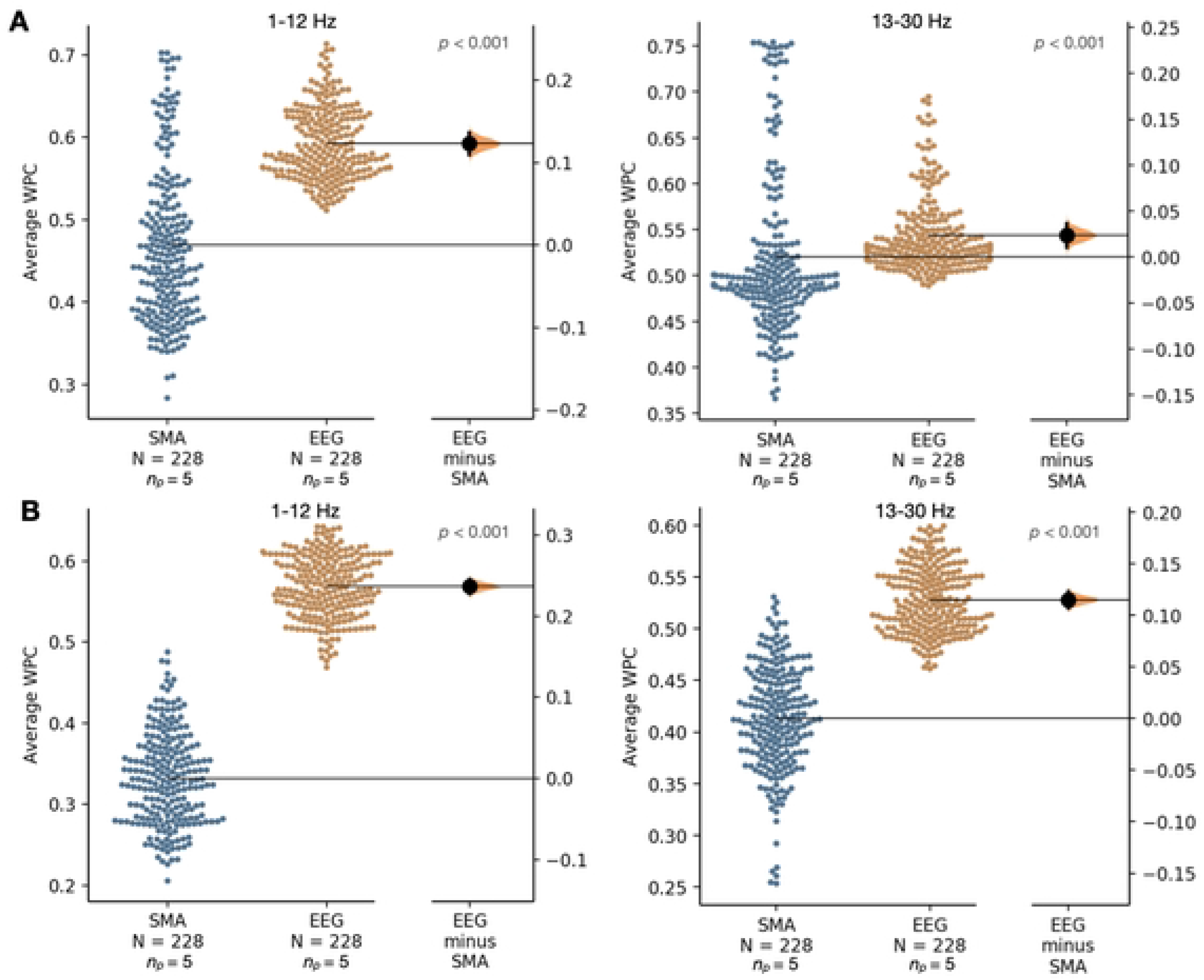
Comparing the average wavelet phase coherence (WPC) of scalp EEG-extracted muscle activity (SMA) and retained (following SMA removal) EEG networks from entire ictal recordings on a per electrode basis (19 electrodes used per recording) for two frequency ranges: 1-12 Hz and 13-30 Hz. (A) For 5 SUDEP patients. (B) For 5 non-SUDEP patients.

### Comparing Average WPC

The average WPC was computed on a per-electrode and per-seizure basis for each of the 13 SUDEP seizures and each of the 12 non-SUDEP seizures. Comparing non-SUDEP to SUDEP patients, the average WPC was significantly higher for SUDEP patients for each of the frequency ranges, as shown in Fig. 6 and 7.

**Figure 6.**
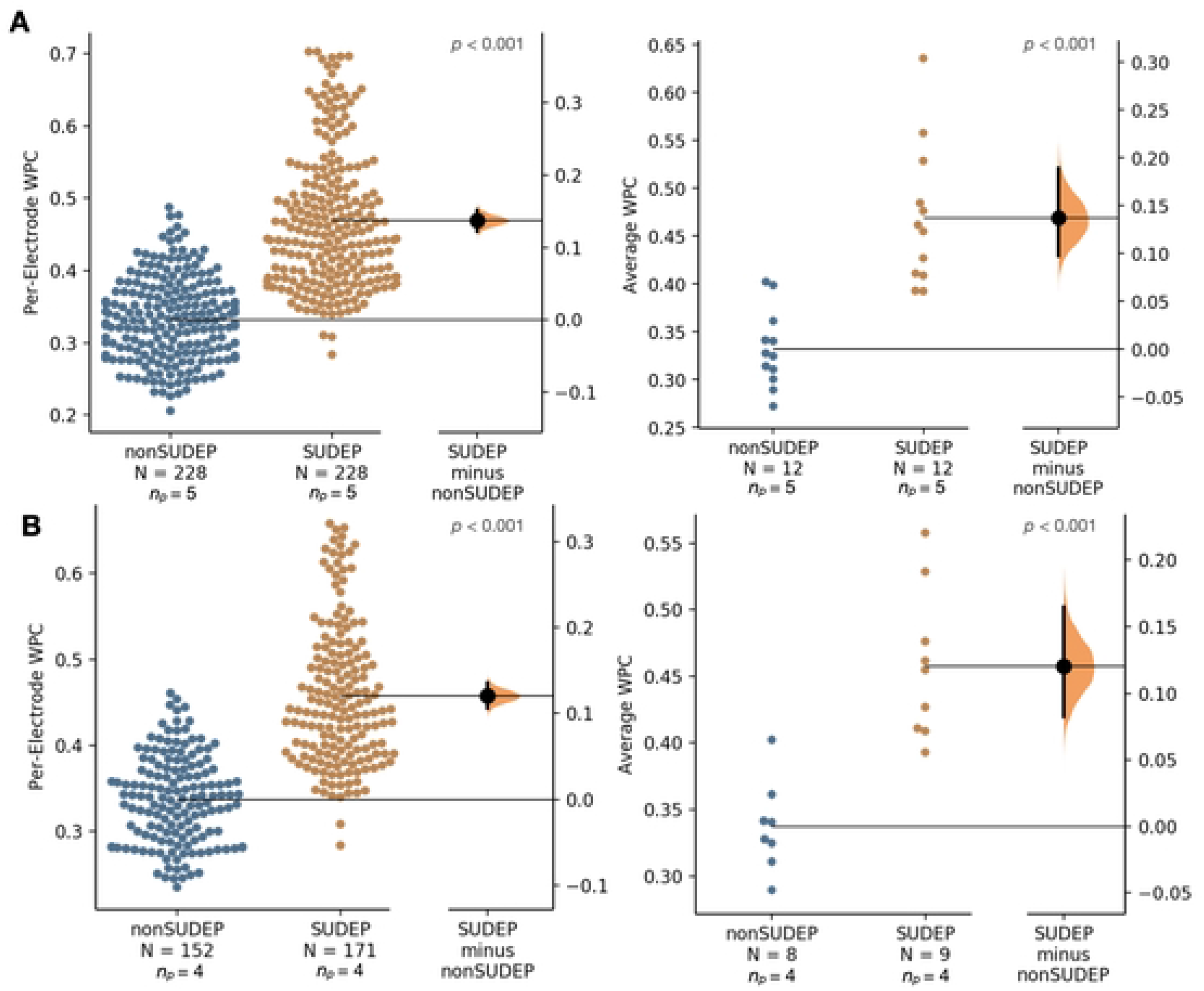
(A) Comparing the average wavelet phase coherence (WPC) over 1-12 Hz for scalp EEG-extracted muscle activity (SMA) networks from entire ictal recordings for 5 SUDEP patients (12 seizures) and 5 non-SUDEP patients (12 seizures) on a per electrode basis (19 electrodes used per recording), left, and mean of all electrodes per seizure, right. (B) Controlling for GTC seizures only, comparing the average WPC over 1-12 Hz for SMA networks from entire ictal recordings for 4 SUDEP patients (9 GTCS) and 4 non-SUDEP patients (8 GTCS).

**Figure 7.**
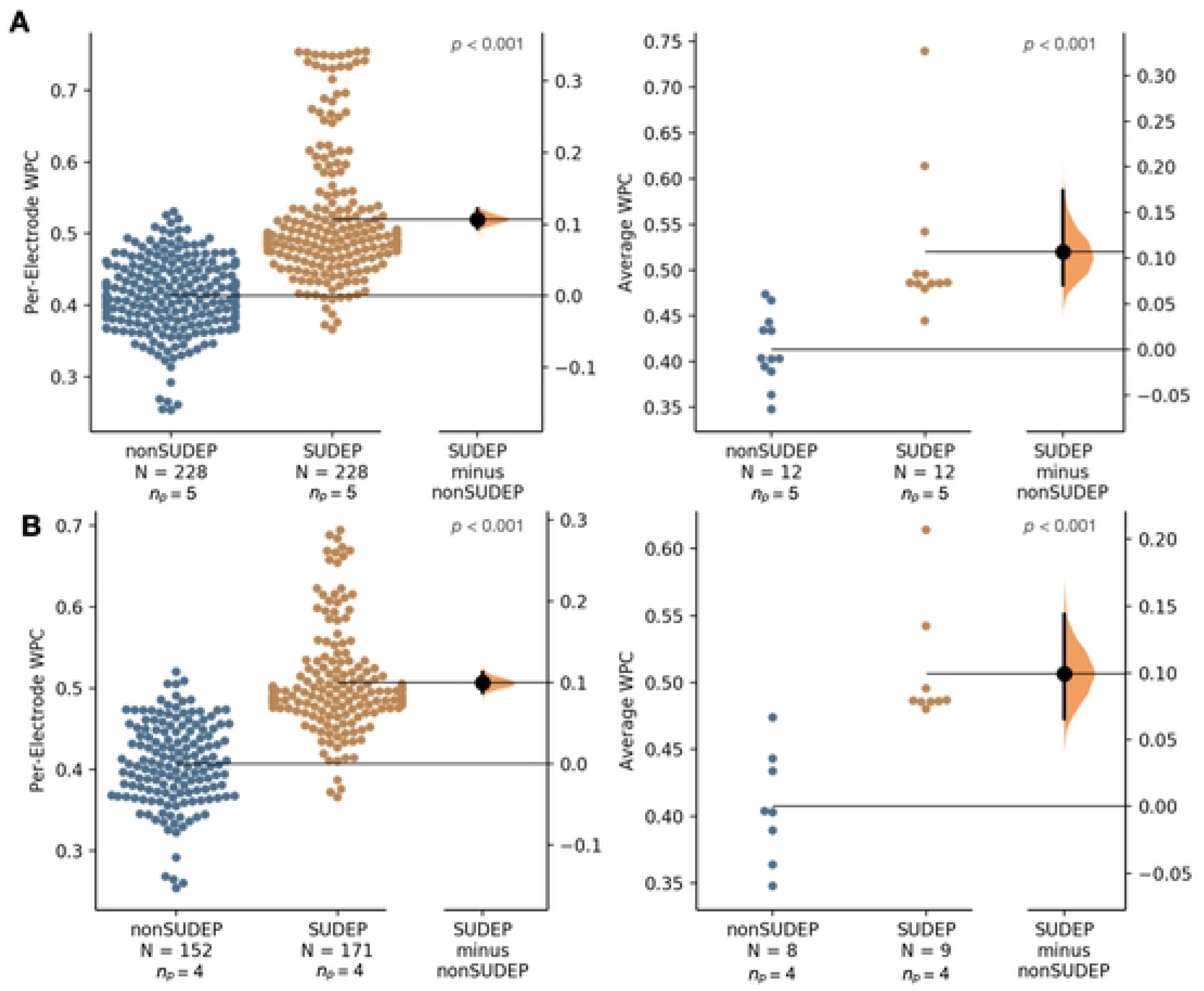
(A) Comparing the average WPC over 13-30 Hz for SMA networks from entire ictal recordings for 5 SUDEP patients (12 seizures) and 5 non-SUDEP patients (12 seizures) on a per electrode basis (19 electrodes used per recording), left, and mean of all electrodes per seizure, right. (B) Controlling for GTC seizures only, comparing the average WPC over 13-30 Hz for SMA networks from entire ictal recordings for 4 SUDEP patients (9 GTCS) and 4 non-SUDEP patients (8 GTCS).

**Figure 8.**
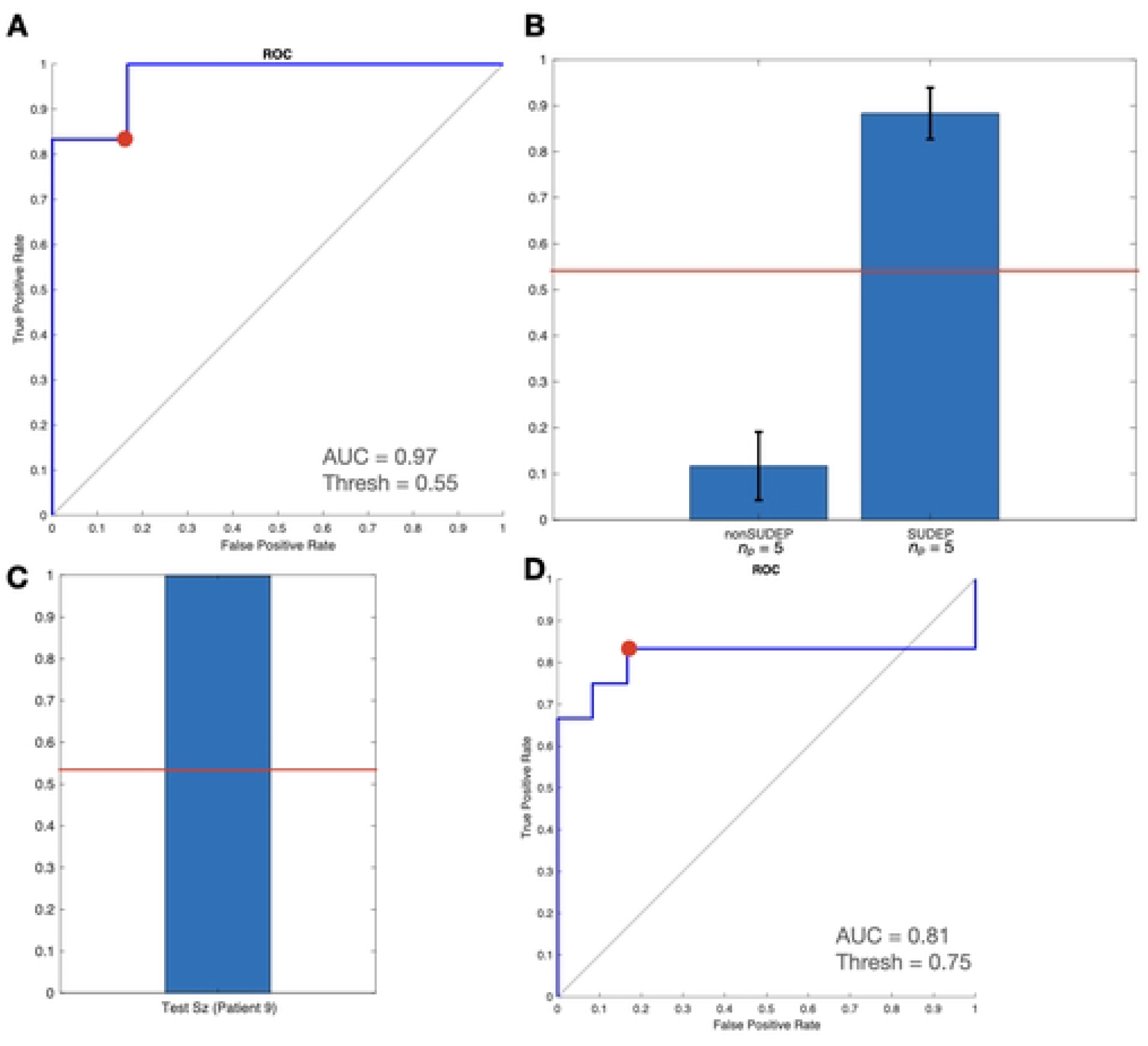
Logistic classifier trained on 1-12 Hz and 13-30 Hz WPC values. (A) ROC curve of training data: Accuracy optimized point (red), used to determine classifier decision threshold. (B) Mean seizure propensity scores, with standard error shown. (C) Propensity score of seizure not included in training set from SUDEP patient. (D) ROC curve for leave-one-out cross-validation, over 10 rounds.

### Risk Assessment

The logistic classifier resulted in a Receiver Operating Characteristic (ROC) curve with an Area Under Curve (AUC) of 97% for training data. The SUDEP seizure withheld for testing was correctly classified as SUDEP by this method.

## Discussion

We found that average WPC was significantly higher for SUDEP compared to non-SUDEP patients for both frequency ranges. Average WPC of SMA is a measure of scalp muscle coherence, as strong contractions would be captured by more electrodes and result in a higher average WPC value. This is in line with previous studies using EMG-EMG coherence in myoclonus assessment in epileptic patients. The possibility that stronger contractions observed in SUDEP patients were due only to the propensity of SUDEP patients to have generalized tonic-clonic seizures [9] was negated as significant differences held in both bands when comparing GTCS only. Further, the risk assessment based on logistic classification of resulting WPC values demonstrated the clinical application of these measures.

This study compared WPC of EMG-like SMA from ictal EEG recordings, as EMG recordings of SUDEP patients were not available. Epilepsy patients treated in an EMU are typically recorded for EEG and ECG; less often for EMG. High-quality ictal EMU recordings of SUDEP patients remain rare; EMG recordings are rarely available for study. Our technique establishes a pathway to using SMA extracted from ictal scalp EEG recordings to leverage EMG-related biomarkers of SUDEP.

This work expands on previous studies [14] extracting EMG-like SMA from EEG recordings. When extracting SMA, there was minimal loss in scalp muscle activity and minimal contamination of EEG signal and noise during the extraction to ensure SMA signals were not distorted or contaminated. SSA was selected as its decomposition process was able to separate the scalp muscle activity signal components from the EEG signal components, in contrast to similar algorithms which were investigated such as ensemble empirical mode decomposition.

Differences in muscular contractions may result from brainstem network disruption implicated in SUDEP. Hypothetical models in rats have suggested that convulsions can result directly from self-sustained epileptic activation in brainstem structures [15], and that these convulsions differ from those originating from the motor cortex. The significantly stronger WPC between extracted SMA in this study may be attributable to convulsions driven by the reticular core of near-SUDEP brainstems.

EMG analysis can detect tonic-clonic seizures in isolation [7, 9] and in multimodal sensory environments [16-17]. Several methods for classification of EMG features have been reported. Empirical mode decomposition of EMG has been used to classify upper limb movements [18], while techniques based on discrete wavelet transforms of EMG have identified muscle movements [19-21].

Video-EEG remains the clinical gold standard for identifying seizures leading to a scarcity of EMG recordings in SUDEP patients. Therefore, it is important to extract EMG features from other modalities. Using only EEG recordings, [22] applied principal component analysis, and both linear discriminant analysis and support vector machines to identify jaw movement, without explicitly identifying EMG. Reference [23] developed an automated system to identify seizures based on ‘optical flow’ of recorded motion. In this study, EMG features were extracted from scalp EEG using SSA.

An EMG-based SUDEP biomarker has been proposed, observing that EMG-derived respiration features identified ictal laryngospasms in mouse models [14]. This possibility was reinforced by a case report of a near-SUDEP patient consistent with this pattern [24].

Investigations of high frequency oscillations (HFOs) in EEG of patients with epilepsy revealed that they were typically of low amplitude and a phase-based measure such as WPC was required for their analysis. Previous work from our team demonstrated that WPC applied to intracranial EEG recordings helped characterize HFOs (80-400 Hz), across brain sites in patients with extratemporal lobe epilepsy that localized seizure onset sites [25]. Subsequent work by the same authors suggested strong coherence between HFO sites in the ictal state, and also in low frequency oscillations (LFOs), 5-12 Hz sites in the interictal state, can localize the same seizure onset sites [26]. Our team also previously reported differences in EEG WPC during infantile epileptic spasms [27].

## Conclusion

SSA with an autocorrelation threshold was an effective method of extracting SMA when using the novel threshold tuning technique mentioned in this paper. The results show that average WPC of ictal SMA is a biomarker for SUDEP. Future research should consider using additional seizure data containing corresponding EMG recordings to help establish a more robust threshold for differentiating scalp muscle activity from brain activity and evaluating additional SUDEP and non-SUDEP ictal SMA data.

## Acknowledgements

B.L. Bardakjian would like to acknowledge support by the Epilepsy Research Program (EpLink) of the Ontario Brain Institute (OBI), the Natural Sciences and Engineering Research Council of Canada (NSERC), and the SciNet HPC Consortium funded by the Canada Foundation for Innovation; the Government of Ontario; Ontario Research Fund (Research Excellence); and the University of Toronto.

